# A new technique for use in the study of the microbiome: An evaluation of a three-dimensional cell culture technique in maintaining the gastrointestinal microbiome of four Balb/c female mice and implications for future studies

**DOI:** 10.1101/734145

**Authors:** Everest Uriel Castaneda, Jeff Brady, Janice Speshock

**Affiliations:** Texas A&M AgriLife Research and Extension Center, Stephenville, Texas, United States of America; Department of Biological Sciences, Tarleton State University, Stephenville, Texas, United States of America

## Abstract

Fluctuations in oxygen, pH, nutrients, or other factors such as food or pharmaceuticals, may perturb the microbiota of the gastrointestinal (GI) tract. This environmental variation is a cause for concern given dysbiosis of the microbiome is correlated with disease states; thereby, model organisms are utilized to study microbial communities during, after, or before shifts in microbes since intact *ex vivo* microbiomes have historically been challenging to utilize. The objective of this study is to culture an explant microbiome of 4 Balb/c, laboratory bred mice to develop an *ex vivo* tool for future microbiome studies. We cultured homogenates of the distal colon of 4 mice in three dimensional, 24 well plate culture dishes. These dishes were incubated for 24 hours in two different oxygen concentration levels, 0% and 20%. The pH of the plate was tested before and after incubation. To analyze the integrity of the microbiome, we utilized 16S sequencing. Further, we utilized 16S metagenomics to characterize fecal samples and colon samples to speculate whether future studies may utilize feces in constructing an explant microbiome to spare animal lives. We found that pH and familial relationship had a profound impact on community structure while oxygen did not have a significant influence. The feces and the colon were similar in community profiles, which lends credence to utilizing feces in future studies. In addition, our efforts successfully cultured archaea, which included difficult to culture strains such as Miscellaneous Crenarchaeota group (MCG) and Methanobacteria. Ultimately, further attempts to culture and preserve an animal’s microbiome needs to control for and maintain stable pH.

## Introduction

The microbiome forms a symbiotic relationship with its host [1]. Essentially, microbes have a cooperative role in the GI tract and contribute to a host’s immune system and metabolism [2–5]. Although the natural relationship between the microbiome and the host is essential, overpopulation by an undesirable species, or dysbiosis, has been linked to particular diseases and phenomena such as autism spectrum disorder [6], cancer [3], and obesity [7]. Historically, researchers have utilized animal models such as gnotobiotic mice to study microbiome-animal interactions but experimentation with such mice is expensive [8–10]. Thus, an *ex-vivo* model of the microbiome that provides cost-effective, reproducible, and reliable results is highly desirable in studying this dynamic [11].

Mouse microbiome studies have expanded our views on the impact of prokaryotes on digestion, disease, and even behavior, but many microbial species cannot persist in culture [12,13]; therefore, most research currently relies on germ-free mice for microbiome studies, which can be cost-prohibitive for many laboratories [7,8,14]. To further assess this transient mixture of microbiota [15], scientists have been utilizing culture independent, next-generation sequencing to inquire about shifts within the microbiome and what stimuli affect these changes in composition [16]. With the decreasing cost of next-generation sequencing, an influx of research has been possible in this area [17]. It is worth noting the financial and ethical burden of raising, sacrificing, and housing model organisms [9,10]; therefore, it would be beneficial to develop techniques to save organisms and further decrease costs.

In this study, we cultured and maintained the GI microbiomes of 4 laboratory-bred female Balb/c mice in three-dimensional (3D) well plates, partitioned into 2 oxygen levels. Due to the variable nature of the oxygen levels of the GI tract [18], we cultured 3D plates in both a conventional incubator and an anaerobic chamber, both at 37 degrees Celsius. Additionally, we determined the microbial composition of the mouse stool and the distal colon to observe if future studies may utilize feces and avoid sacrificing organisms altogether. We utilized sequencing methods to verify final proportional community composition of each sample.

## Materials and Methods

### Subjects

The study was performed under a protocol approved by the Tarleton State University Institutional Animal Care and Usage Committee (Animal Use Protocol 12-009-2016-A1). 4 Balb/c females 8 weeks in age were utilized in this experiment. Females were housed together and raised on identical chow diets and similarly weaned. Mice 1 and 2 were siblings while mice 3 and 4 were siblings. The siblings were born from different dams and sires. All mice were held in sterile containers and euthanized with 150 µL of sodium pentobarbital delivered intraperitoneally. Post injection, mice shed two to three samples of stool which were recovered utilizing sterile forceps and immediately frozen. Once deceased, 2.5 cm of the large distal colon were removed. After, two to three small additional 0.5 cm samples of the large distal colon were excised from the specimen and immediately frozen. Colon tissue extractions were added to a sterile tissue grinder along with 5 mL of Dulbecco’s Modified Eagle Medium (DMEM). The sample was manually homogenized into a liquid solution. Homogenate was checked for pH by applying a small droplet with a mechanical pipette onto litmus paper.

### Culture method

Cultures were established in sterile 24 well plates with multiwell tissue culture inserts, 8 µm pore size (Corning Incorporated, New York). Corning PuraMatrix Peptide Hydrogel was prepared using 8 mL of molecular grade water and 20 ml of hydrogel to create a 0.25% solution. 150 mL of the prepared solution were added to each of the 12 well inserts. In addition, 500 mL of supplementary DMEM was added under each well insert. This technique is a modified version of 3D tissue culture repurposed for culturing our *ex-vivo* microbiome. Once the 3D culture plates were prepared, 250 mL of the homogenized colon were added to each of the prepared 12 well inserts. Plates were checked for baseline pH by transferring a small drop of medium with a mechanical pipette onto litmus paper. The plates were then added to a single incubator, but to create an anoxic environment, half of the plates were incubated in an anaerobic chamber. Plates were incubated for 24 hours. Each insert’s medium was then tested for pH again and transferred into sterile 2.5 mL storage tubes and frozen for future DNA extraction.

### DNA extraction and library production

DNA was extracted from each sample using a solid phase extraction protocol from Brady et al. [19]. After extraction, DNA was amplified utilizing prokaryote-specific primers, 519F 5’-CAGCMGCCGCGGTAA-3’) and 785R (5’-TACNVGGGTATCTAATCC-3’), that target the V4 region of the 16Ss rRNA [20,21]. PCR amplification was accomplished through denaturation at 95°C for 3 minutes, followed by 35 cycles of 95°C for 10 seconds, 55°C for 30 seconds, and 72°C for 30 seconds. Dual 6 bp DNA barcodes were added to sequencing libraries using the same PCR protocol and Illumina P5 and P7 flowcell binding adapters [22]. Sequencing libraries were size-selected with a Pippin Prep instrument (Sage Science, Beverly, MA) to a length of 300-600 base pairs. Sequencing was conducted on a MiSeq instrument using 600 cycle paired end v3 sequencing kits at the Texas A&M University Genomics Core Facility. Raw sequences were processed with QIIME [23] and USEARCH [24]. Taxonomy was assigned using the Greengenes 13.8 database [25] as a reference with UCLUST [24], and reference-based Operational Taxonomic Unit (OTU) picking was conducted at 97% sequence similarity using the RDP [26] method in QIIME.

### Statistical analysis

Cumulative Sum Scaling [27] was utilized to normalize the data. Biom files were constructed with QIIME [28] and transferred into R [29] and Microsoft Excel for further statistical analysis. Phyloseq [30], ggplot2 [31], and vegan [32] packages were utilized to evaluate alpha and beta diversity with seed set at 1400. Alpha diversity was assessed using the Shannon diversity index. Variation in alpha diversity for oxygen, pH, mouse, feces, and colon comparisons were first checked for normality using the Shapiro-Wilk test for normality [33]. The data was non-normal in distribution (Shapiro-Wilk test, w=0.9506, p<0.01); therefore, comparisons were made with non-parametric tests. All multivariate tests were corrected using false discovery rate (FDR) [29]. Comparisons of alpha diversity were conducted using Kruskal Wallis one-way analysis of variance (KW ANOVA) and Wilcoxon rank sums test (Wilcoxon test) while comparisons of beta diversity were assessed with unweighted unifrac distance metrics at 1000 permutations using permutational multivariate analysis of variance (PERMANOVA). Dunn’s test, post-hoc analysis was conducted using the dunn.test package in R [34]. In addition, non-parametric t-tests were used for comparisons of mean abundance in individual bacterial strains between samples. Principle coordinate analysis (PCoA) and canonical correspondence analysis (CCA) were performed at the level of OTU using unweighted unifrac distance metrics. PRIMER 7 [35] was used for the hypothesis testing utilizing square root transformed Bray-Curtis ordination data at 9999 permutations. A microbial network was constructed using the Co-occurrence Network Interferences (CoNet) [36] application for Cytoscape [37]. Feces and colon data were removed before CoNet analysis. CoNet has been utilized in previous studies to investigate defined interactions between microbes [38–40]. Spearman correlation coefficient with a cutoff ratio of 0.6 was utilized, and to focus the network, only microbes with sequence counts greater or equal to 20 were included. 1,000 permutations were accomplished through a bootstrapping method with an FDR correction [39].

## Results

### pH and oxygen

pH readings of each plate were taken before and after incubation. As shown in Table 1, pH fluctuated from the original homogenate and baseline (before 24-hour incubation). In addition, mouse samples maintained varying levels of pH which correlated with differences in oxygen concentration (Table 1).

**Table 1.** Sample size, pH, and oxygen level.

| Mouse | Homogenate pH | Sample Size | Oxygen Level | Plate Baseline pH | Plate Final pH |
| --- | --- | --- | --- | --- | --- |
| M1 | 7.5 | 12 | 20% | 8 | 10 |
|  |  | 11 | 0% | 8 | 9 |
| M2 | 7.5 | 11 | 20% | 8 | 10 |
|  |  | 12 | 0% | 8 | 9 |
| M3 | 7.5 | 12 | 20% | 8 | 7 |
|  |  | 11 | 0% | 8 | 6 |
| M4 | 7.5 | 12 | 20% | 8 | 7 |
|  |  | 12 | 0% | 8 | 6 |

### 16s rRNA sequencing results

We analyzed microbiomes from the 4 mice at varying oxygen levels after culturing them *in vitro*. In the 24 well plate system, we utilized 12 membrane inserts to culture the microbiomes. After mice succumbed to euthanasia, colon samples were harvested to complement fecal shedding. Mouse 1, 2, and 4 shed two fecal samples each while mouse 3 shed three fecal samples; therefore, we had a total of 9 colon and 9 fecal samples across 4 mice. After sequence quality filtering we had a total sample size of 111 samples and 3,133,666 sequences total (Table 2).

**Table 2.**
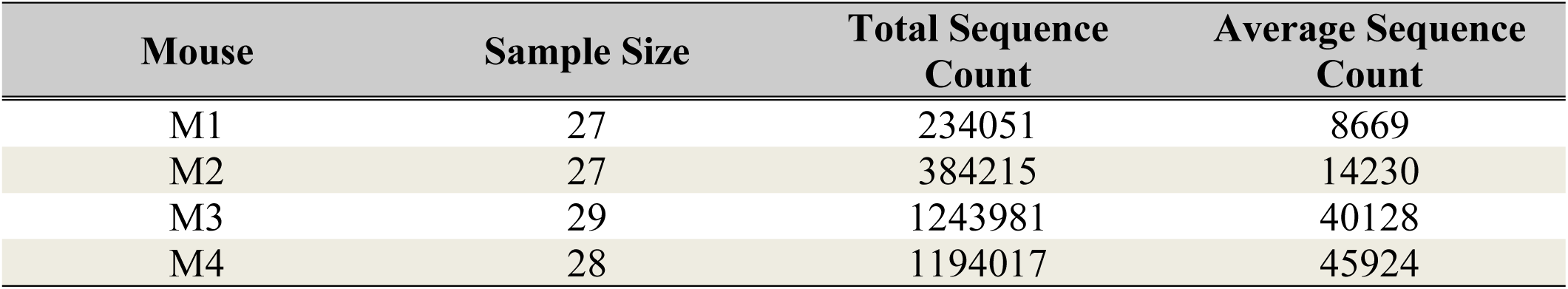
Total sample size and sequence count per mouse.

### Fecal and colon comparison

The feces and the colon samples were characterized for microbial composition at the phylum and family level (Figs 1A and 1B, respectively). Community composition was dominated by the phyla Firmicutes and Bacteroidetes (Fig 1A), with means of 47% and 49%, respectively, and standard deviations (SD) of 23%. In addition, the *Bacteroidales* family S24-7 was highly abundant with a mean of 42% and a SD of 20% (Fig 1B). These results are consistent with recent microbiome studies of mice [41–43]. Beta diversity analysis for each of the feces and colon samples showed no difference in composition (Table 3), and alpha diversity analysis (Fig 1C) also revealed no difference (KW ANOVA, chi-squared = 6.64, df = 7, p = 0.47). Therefore, all colon samples were pooled together, and all feces samples were pooled together for a statistical comparison of feces and colon. The Shannon diversity index was utilized for comparison of the bulk samples (Fig 1D). Results showed no difference between the feces and colon (Wilcoxon test, p = 0.44). In addition, beta diversity comparison showed no difference (PERMANOVA, Pseudo F = 1.06, p = 0.37). Since we found that feces and colon samples are similar, we pooled sequences from all feces and colon samples together into one bulk sample, named “microbiome,” for diversity comparisons with cultured microbiomes.

**Table 3.**
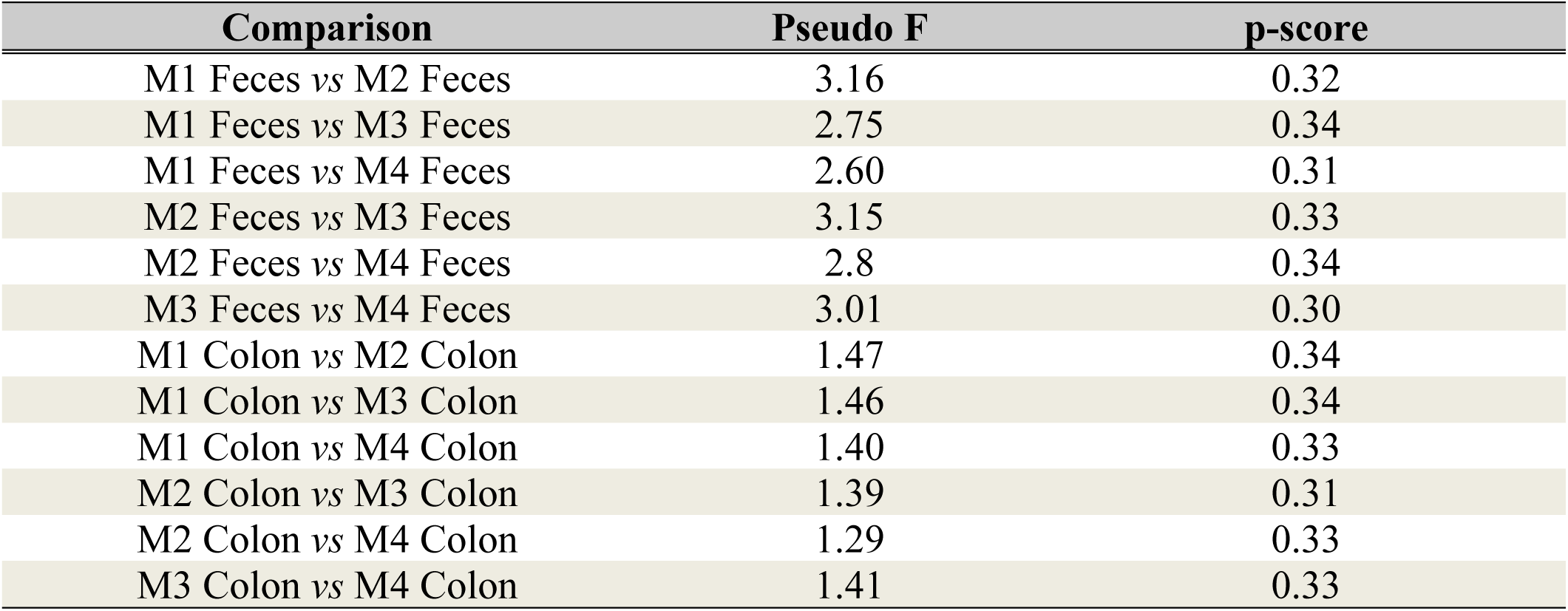
Results of the pairwise PERMANOVA tests between feces and colon samples.

**Fig 1.**
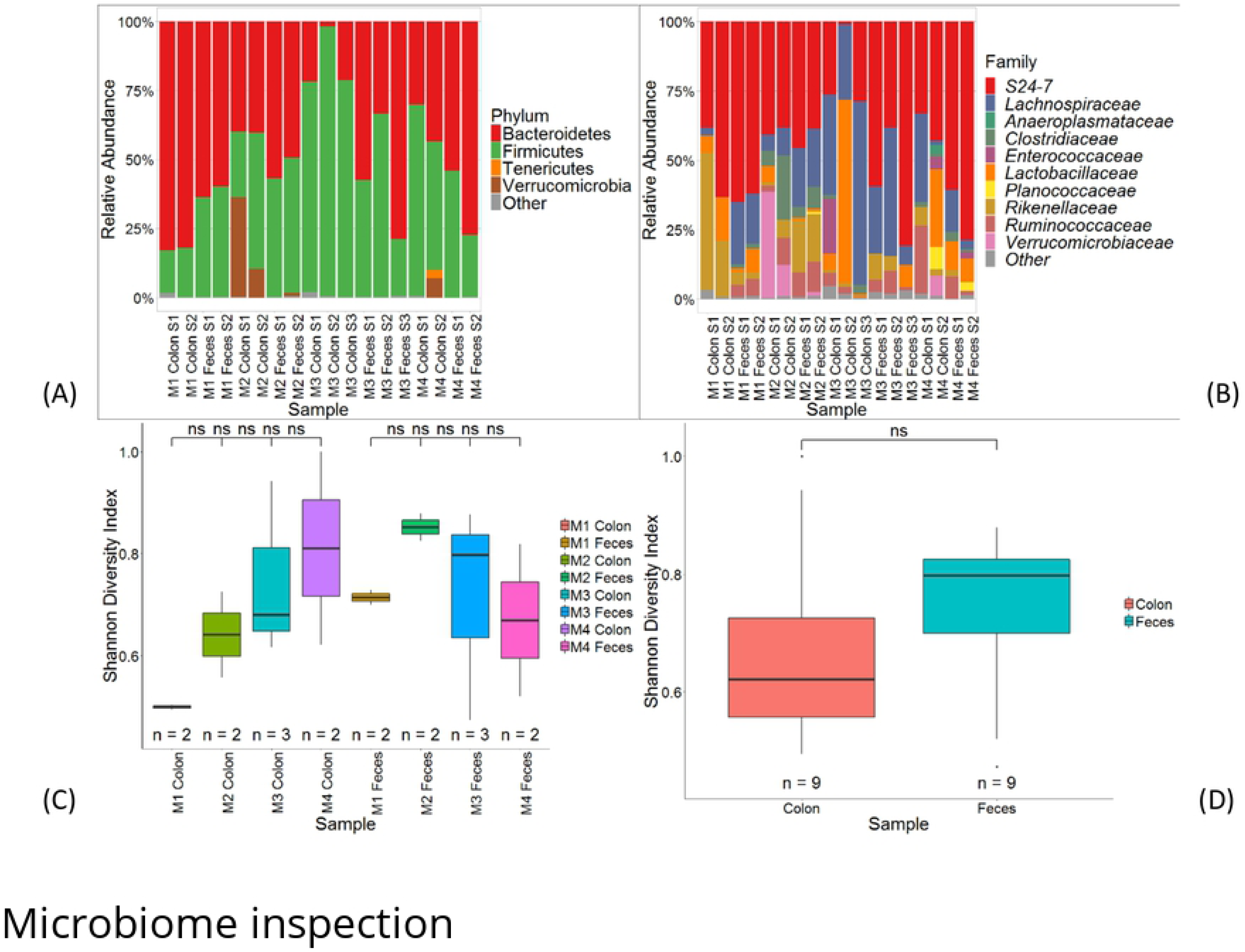
Community composition of fecal and colon samples. (A) Relative abundance of bacterial phyla. Phyla with observations less than 1% are pooled into “Other” category. The first two characters represent the mouse from which the organ and stool were dissected. “S” denotes the different samples acquired. (B) Relative abundance of bacterial families. Families with observations less than 1% are pooled into “Other” category (C) Comparison of Shannon diversity between feces and colon samples. “ns” means non-significant, p>0.05. (D) Shannon diversity comparison of pooled feces and colon samples.

### Microbiome comparison

In the explanted microbiomes, Firmicutes and Bacteroidetes were the dominant phyla (Fig 2A). Firmicutes had the highest average relative abundance, 70% (SD 28%), with Bacteroidetes averaging 18% (SD 16%). Across all cultures, *Enterococcus* was highly abundant (Fig 2B) having a mean of 47% (SD 32%). The Shannon diversity index was different between the explanted cultures and the microbiome of the mice (KW ANOVA, chi-squared = 73.58, df = 8, p < 0.01). Post-hoc analysis shows that, compared to the microbiome, mouse cultures 1 and 2 were the same while mouse cultures 3 and 4 differed (Fig 2C). The microbial profile of each sample revealed a difference in beta diversity between the plates and the microbiome (Table 4), which is also reflected in the PCoA plot (Fig 2D).

**Table 4.** Results of the pair-wise PERMANOVA tests between cultured plates and harvested samples.

| Comparison | Pseudo F | p-score |
| --- | --- | --- |
| M1 0% vs Microbiome | 11.09 | $p < 0.01$ |
| M1 20% vs Microbiome | 18 | $p < 0.01$ |
| M2 0% vs Microbiome | 11.93 | $p < 0.01$ |
| M2 20% vs Microbiome | 15.21 | $p < 0.01$ |
| M3 0% vs Microbiome | 14.94 | $p < 0.01$ |
| M3 20% vs Microbiome | 7.95 | $p < 0.01$ |
| M4 0% vs Microbiome | 18.83 | $p < 0.01$ |
| M4 20% vs Microbiome | 21.321 | $p < 0.01$ |

**Fig 2.**
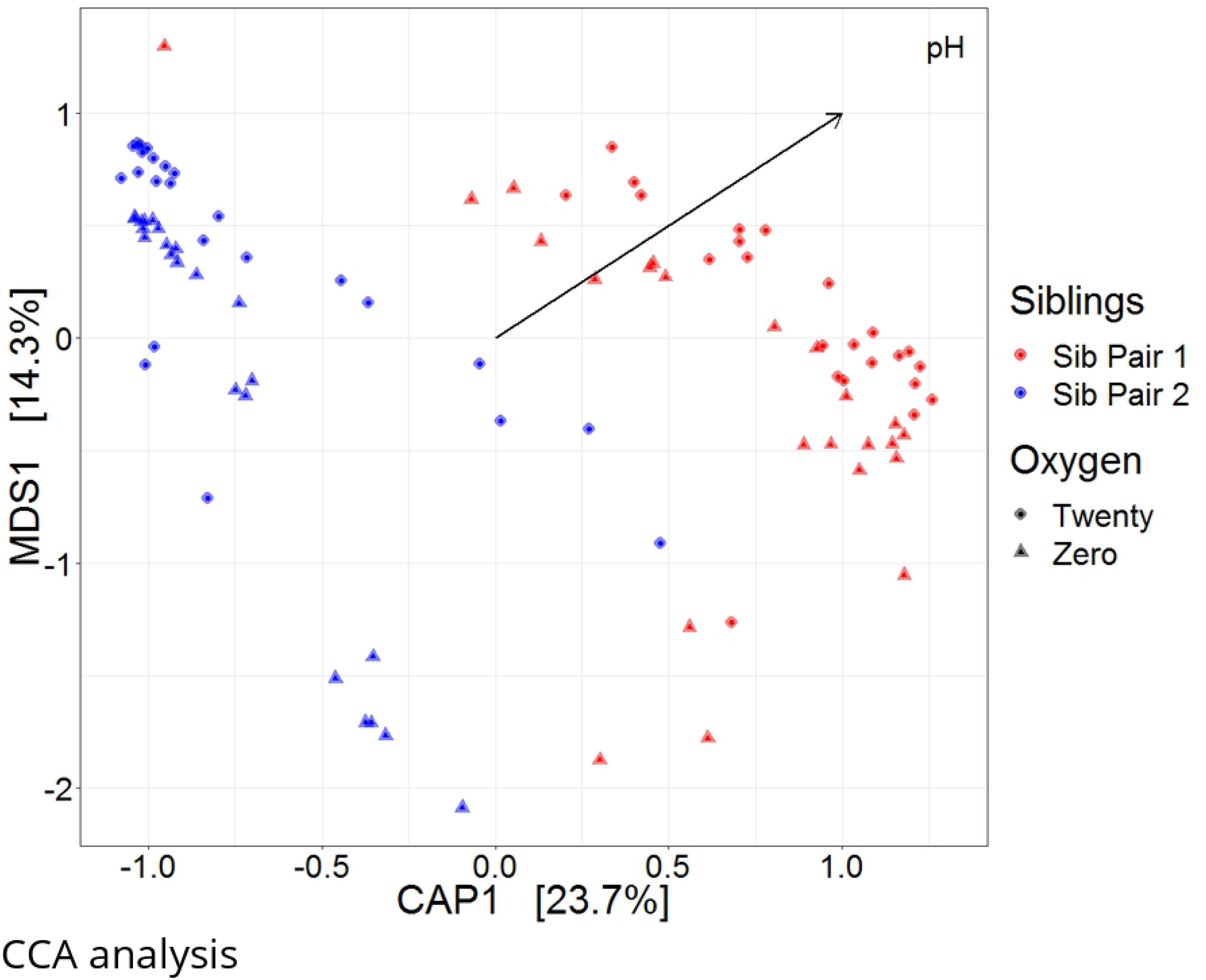
Community composition of cultures and comparison of microbiome. (A) Relative abundance between bacteria Phyla. Phyla with observations less than 1% are pooled into “Other” category. The first two characters represent the mouse, and the percent oxygen used in culture conditions is noted. (B) Relative abundance of bacterial genera. Genera with observations less than 5% are pooled into “Other” category. (C) Results of the post-hoc, Shannon Diversity, pairwise comparisons between cultures and the microbiome. p<0.05 is noted by “*”, p<0.0001 is noted by “****”, and non-significance is noted by “ns.” (D) Unweighted unifrac PCoA plot for plates and microbiome comparison. 12 well plates are denoted by the mouse in which they were derived from, “M,” and the percent oxygen.

### Environmental variables

A community profile of cultural composition due to varying levels of oxygen and pH was constructed at the phylum and genus levels (Fig 3A-D). No difference in alpha diversity existed between the two oxygen levels (Wilcoxon test, p = 0.34; Fig 3E). Results showed similarity in beta diversity (PERMANOVA, Pseudo F = 1.25, p = 0.21). Multivariate analysis shows a significant difference in alpha diversity associated with differences in pH (KW ANOVA, chi-squared = 58.13, df = 3, p < 0.01). Post-hoc analysis revealed that plates reaching a pH of 6 and 7 were similar while all other comparisons differed (Fig 3F). Comparisons of microbial communities also showed a marked difference between plates of varying pH levels (Table 5). Additionally, CCA and PERMANOVA revealed that oxygen did not contribute to community clustering (Table 6; Fig 4).

**Table 5.** Results of the pairwise PERMANOVA test between pH groups.

| Comparison | Pseudo F | p-score |
| --- | --- | --- |
| pH 6 vs pH 7 | 1.86 | 0.04 |
| pH 6 vs pH 9 | 17.15 | $p < 0.01$ |
| pH 6 vs pH 10 | 25.46 | $p < 0.01$ |
| pH 7 vs pH 9 | 15.27 | $p < 0.01$ |
| pH 7 vs pH 10 | 22.31 | $p < 0.01$ |
| pH 9 vs pH 10 | 1.62 | 0.03 |

**Table 6.** Hypothesis testing for sources of variation based on PERMANOVA analysis.

| Source | df | SS | MS | Pseudo F | P(perm) | Unique perms |
| --- | --- | --- | --- | --- | --- | --- |
| Oxygen | 1 | 3432.6 | 3432.6 | 1.6677 | 0.0756 | 9903 |
| Sibling Relationship | 1 | 52930 | 52930 | 25.716 | 0.0001 | 9921 |
| Oxygen x Sibling Relationship | 1 | 2407.4 | 2407.4 | 1.1696 | 0.2437 | 9915 |
| Residuals | 89 | 1.8319E+05 | 2058.3 |  |  |  |
| Total | 92 | 2.4194E+05 |  |  |  |  |

df, degrees of freedom; SS, sum of squares; MS, mean square.
The only significant effect fitted in the PERMANOVA is sibling relationship (fixed factor) while oxygen (fixed factor) is not significant. Sibling relationship explains the changes in community composition as time elapsed in the incubators.

**Fig 3.**
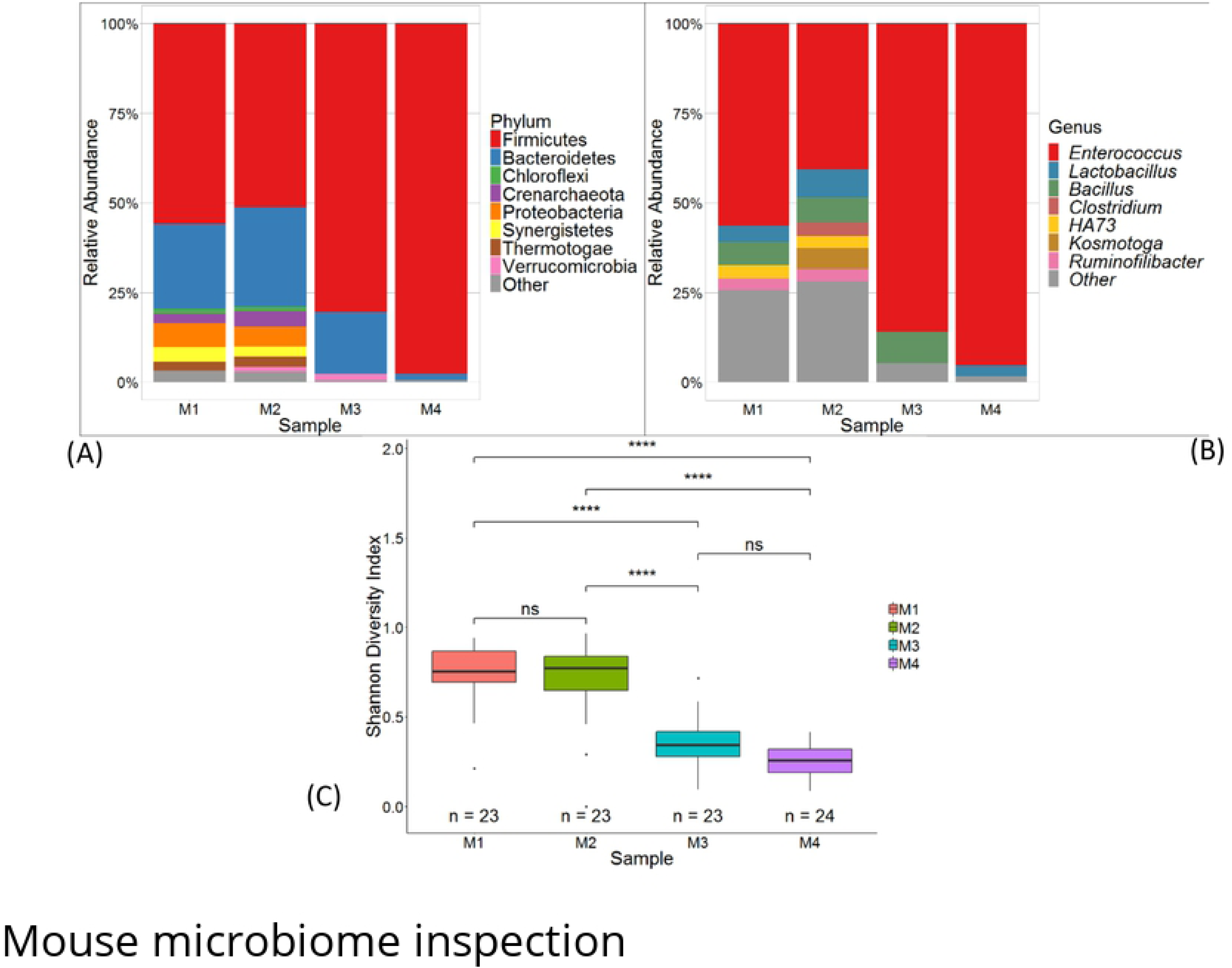
Microbial structuring due to environmental factors. (A) Relative abundance between bacteria phyla. Phyla with observations less than 1% are pooled into “Other” category. (B) Relative abundance of bacterial genera. Genera with observations less than 1% are pooled into “Other” category. (C) Shannon diversity comparison between oxygen concentrations. Non-significance is shown by “ns.”

**Fig 4.**
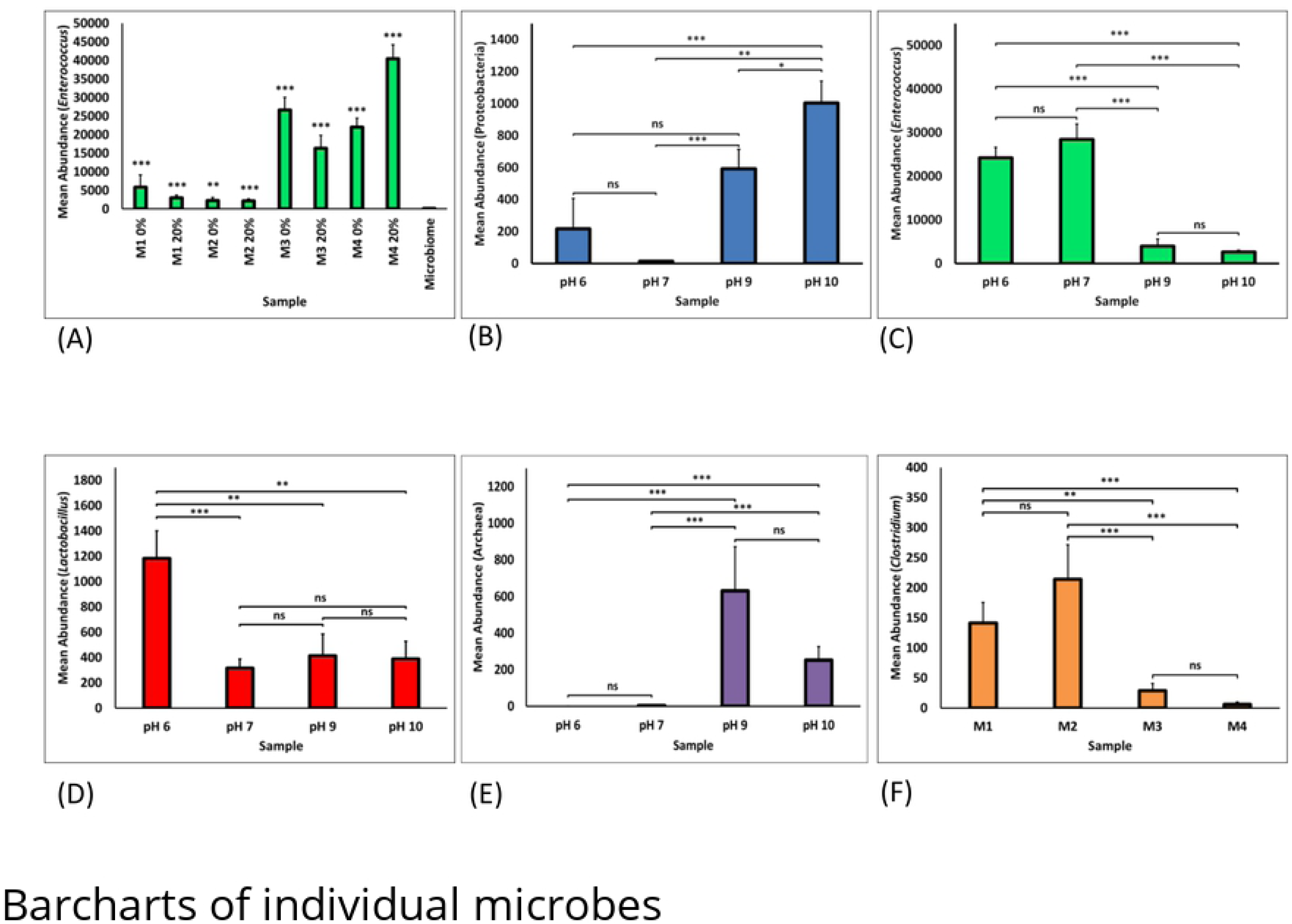
CCA for the effects of pH, oxygen, and sibling relationship on community structuring.

### Siblings

A marked difference in cultural composition was noted by familial relationship (Table 6, Fig 4, Fig 5A, B). Not only was clustering associated with siblings, shown in Fig 4, but Shannon diversity significantly varied (KW ANOVA, chi-squared = 56.24, df = 3, p < 0.01). Post-hoc analysis showed mouse 1 and mouse 2 were similar and varied from mouse 3 and mouse 4, which were also similar (Fig 5C). Additionally, beta diversity varied according to familial relationship (PERMANOVA, Pseudo F = 12.82, p < 0.01).

**Fig 5.**
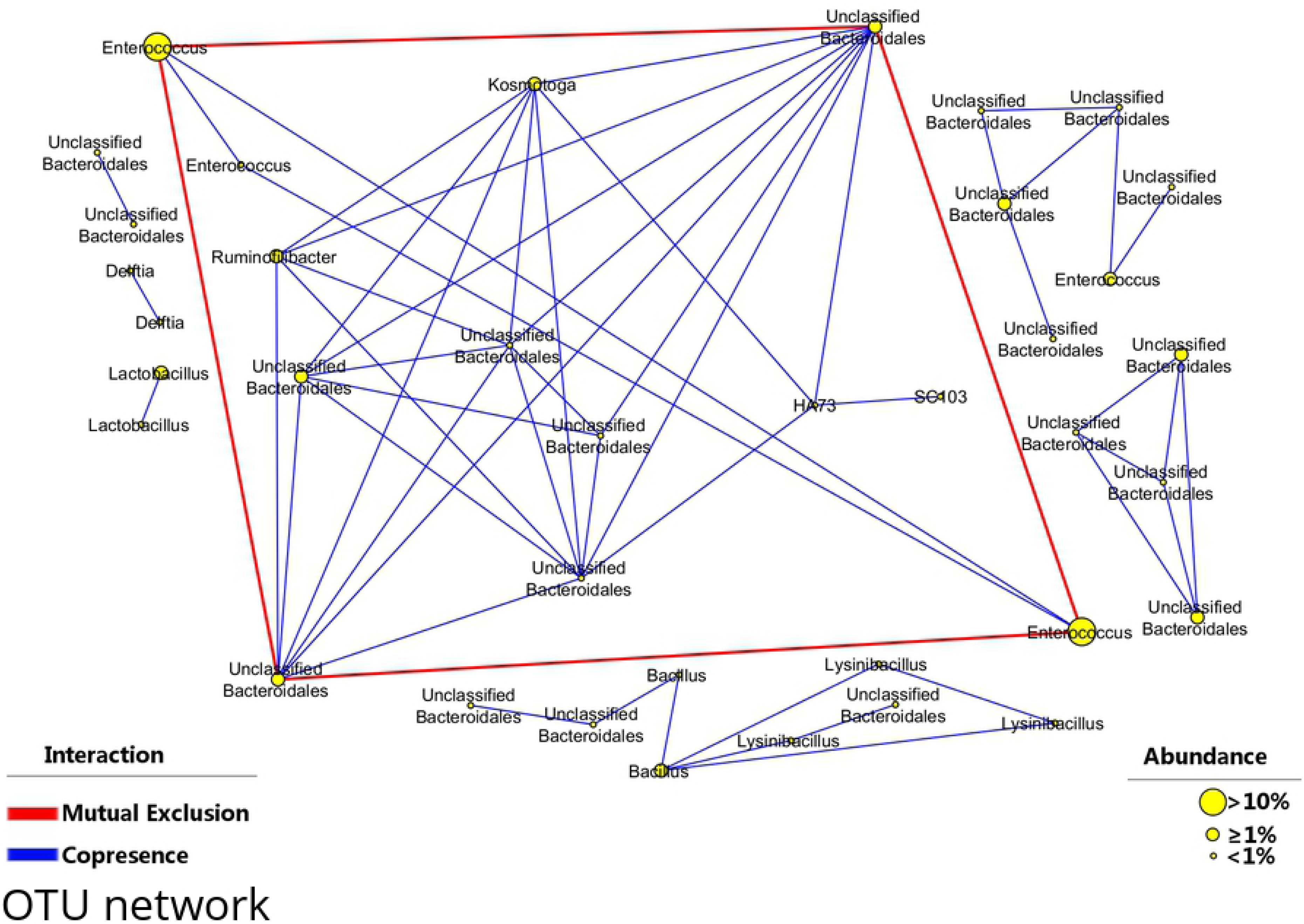
Microbial composition by mouse. (A) Relative abundance between bacteria phyla. Phyla with observations less than 1% are pooled into “Other” category. (B) Relative abundance at the level of genus. Observations less than 0.03% are pooled into “Other” category. (C) Shannon diversity index comparison using mouse 2 as a reference group. “****” means significance p<0.0001 while “ns” means non-significant.

### Individual Strain Comparisons

*Enterococcus* significantly increased between mice cultures and the microbiome (Fig 6A). Additionally, Proteobacteria strains were more abundant in cultures reaching a high pH; although, cultures reaching pH 6 were equivalent in Proteobacteria compared to cultures reaching pH 9 (Fig 6B). *Enterococcus* strains also followed a similar dynamic in which they increased in cultures reaching a lower pH, 6 and 7 (Fig 6C). Subsequently, cultures with the lowest pH, 6, had a significantly high abundance of *Lactobacillus* (Fig 6D). Archaea were more abundant in plates with pH 9 and 10 (Fig 6E), and further, *Clostridium* strains were more likely to be present in mouse 1 and 2 cultures compared to mouse 3 and 4 (Fig 6F).

**Fig 6.**
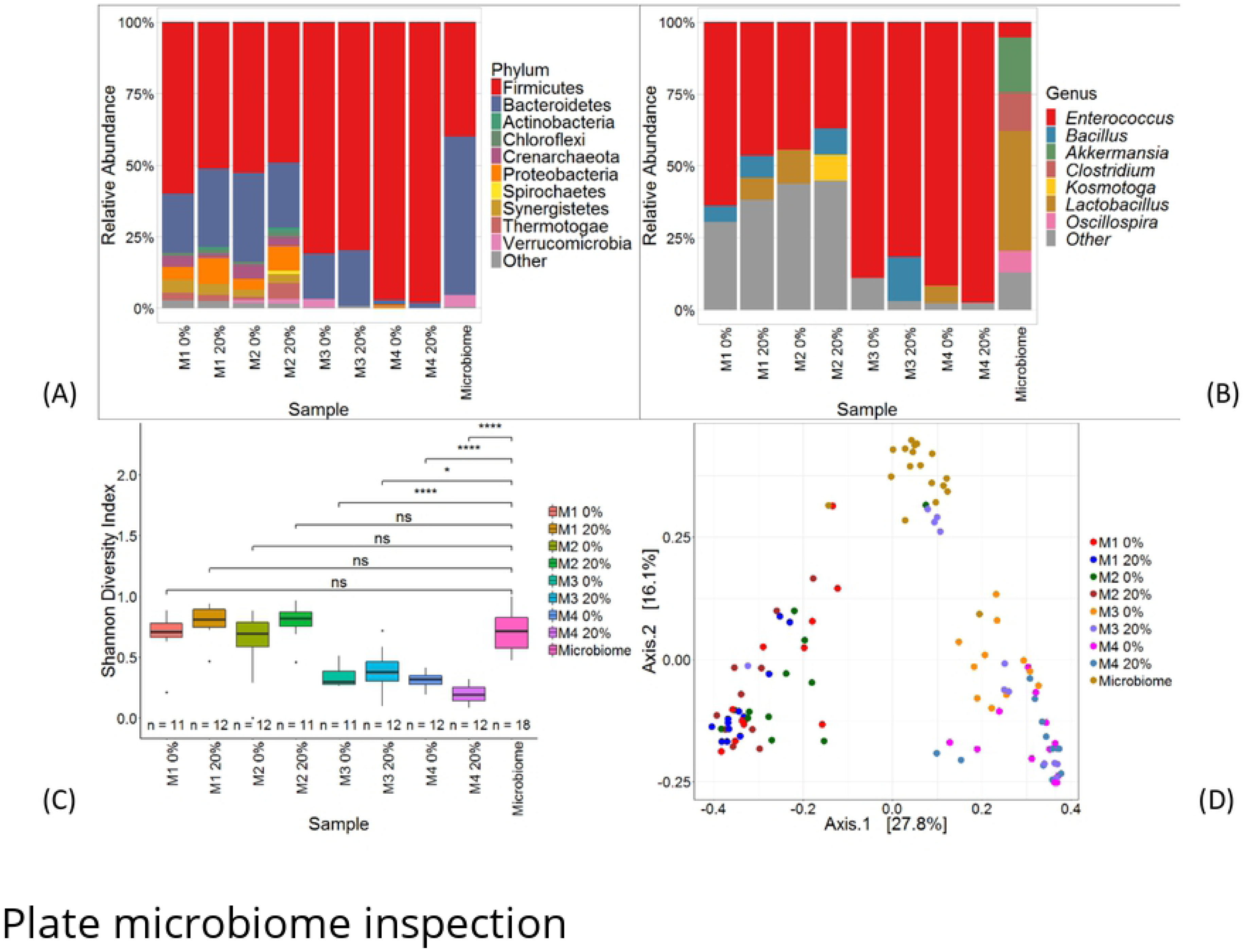
Comparison barcharts of individual microbial taxa. (A) Comparison of the mean abundance of *Enterococcus* in cultured plates compared to the microbiome. (B) Comparison of the mean abundance of Proteobacteria in samples with varying pH. (C) Comparison of mean abundance of *Enterococcus* in samples with varying pH. (D) Comparison of mean abundance of *Lactobacillus* in samples with varying pH. (E) Comparison in mean abundance of archaea in samples with varying pH. (F) Comparison in mean abundance of *Clostridium* in cultures partitioned by which mouse, M, it originated. “ns”, non significant; “***”, p < 0.001; “**”, p < 0.01; “*”, p ≤ 0.05.

### Microbial network

The OTUs in the microbial network represent 88% of the relative sequence count for the cultured well plates (Fig 7). Much of the interactions were positive in nature meaning copresence in a shared-niche is the most abundant interaction type. Negative, mutually exclusive interactions are only between OTUs from the genus *Enterococcus* and several OTUs from the order Bacteroidales (Fig 7). The interaction between the 4 mutually exclusive OTUs account for 50% of all sequences.

**Fig 7.**
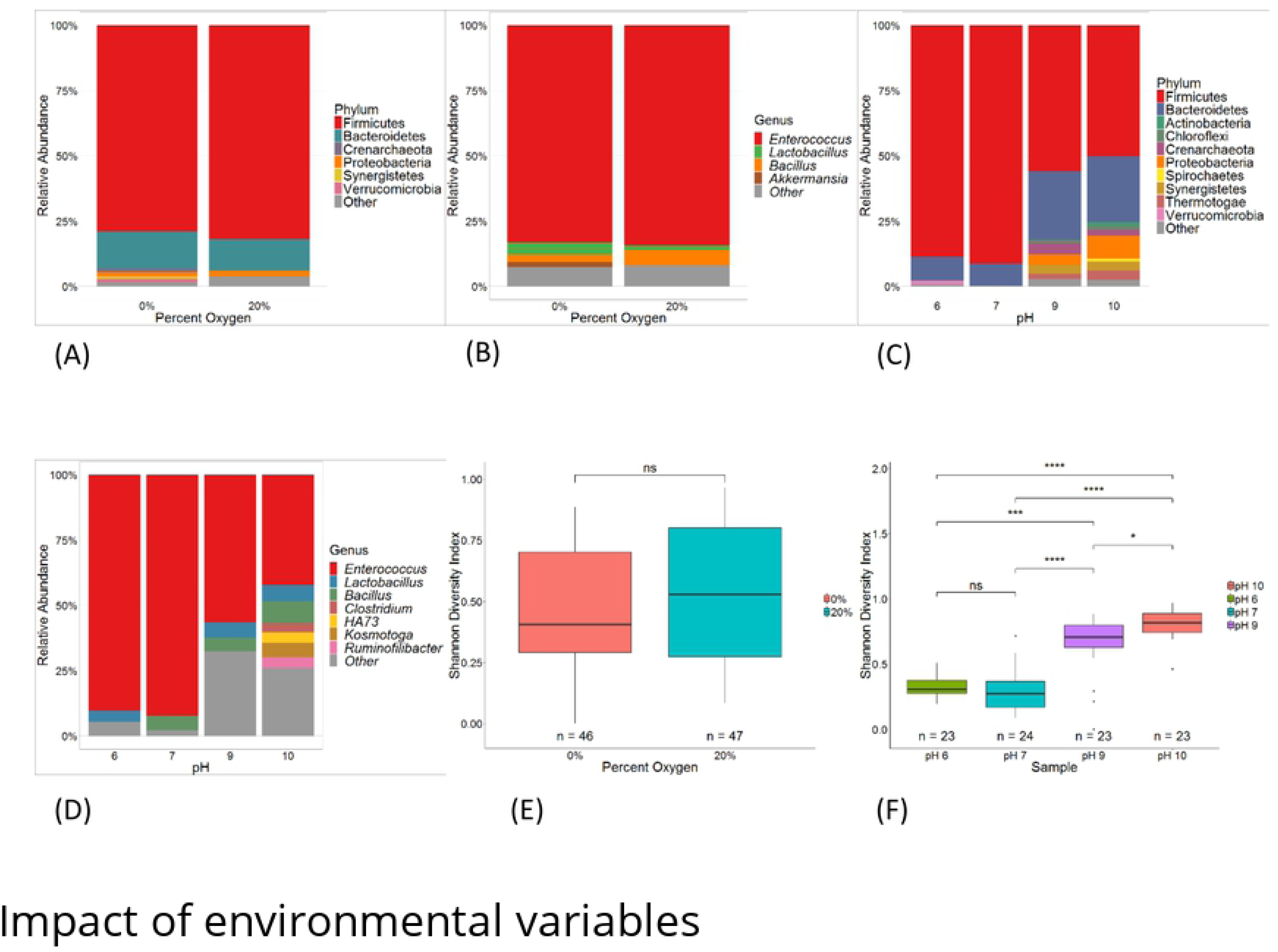
Microbial network generated using Spearman’s rank correlation at the taxonomic level of genus. Most of the edges, 54 of 58, are of a positive correlation. The rest are negative. Nodes sizes are configured based on abundance. Nodes greater than or equal 10% are the largest, intermediate sized nodes are greater than or equal to 1%, and the smallest nodes are less than 1%.

## Discussion

Many studies of the microbiome utilize germ-free mice, which are expensive to house and breed and require sacrificing animals to study the GI microbiome [14,16]. In this study, we attempted to culture an *ex vivo* microbiome in 3D well plates to decrease the cost associated with studying animal microbiomes. We found that cultures for mouse 1 and 2 were comparable to the gut microbiome in Shannon diversity (Fig 2C), which is very promising. Ultimately, our explant microbiome was significantly different than the *in vivo* microbiome, but we were able to culture a diverse number of prokaryotic strains utilizing our method. Optimizing efforts in culture media, detection, and atmospheric gradients is extremely important in culturing desired microbes [44]. With very little optimization, we were able to culture many gut microbial species including difficult to culture strains such as Methanobacteria [45], mean of 0.57 (SD 4.31), and MCG [46], mean of 218.59 (SD 633.71), which included the fecal B10 strain [47], (Fig 6E).

The upsurge of *Enterococcus* in cultured plates (Fig 6A) is explained by its competitiveness outlined in Fig 7. A likely scenario is that *Enterococcus* outcompeted strains within the order Bacteroidales, which makes up a high proportion of the gut microbiome [48,49]. *Enterococcus* is a facultative anaerobe [50] that may respire aerobically in the presence of hemin using cytochrome *bd* terminal oxidase, which reduces oxygen into water [51,52]. Since it is a known pioneer colonizer of the GI tract, its presence possibly established and maintained the anoxic environment by metabolically depleting the atmospheric oxygen, which allowed obligate anaerobes to thrive [53,54].

Post-incubation, a shift in pH was seen amongst the various 12 wells of the plates (Table 1). This shift in pH is accounted for by the increase in gram negative Proteobacteria in plates with a more basic pH (Fig 6B). Creation of the amine groups from this phylum perhaps led to the increase in pH [55–57]. Further, the decrease in pH may be related to the increase in the lactic acid fermenter *Enterococcus* in plates with a pH of 7 (Fig 6C) [52] while plates with a pH of 6 may be partially explained by the increase in both *Enterococcus* and *Lactobacillus* (Fig 6C, D) [52,58,59]. Additionally, pH was a strong influencer in the growth of archaea. Archaea grew more readily in plates with a higher pH (Fig 6E). Not only are archaea difficult to culture, but also their diversity is not well studied in regard to the gut microbiome [60]. Results are comparable to those of Ilhan et al. wherein pH had a strong influence on microbial composition in fecal anaerobic cultures [59]. There was a possible interaction between differences in the microbial communities seeding the culture plates and physio-chemical culture conditions causing the pH in plates from different mice to swing in opposite directions. Additionally, culturing plates in an anaerobic chamber instead of a CO_2_ incubator may have exacerbated pH instability [61]. Future efforts to stabilize pH may allow for additional growth of archaea and provide a means to temporarily culture and study members that have in the past been recalcitrant to culture methods.

The microbiome is passed on from mother to litter [8,62]. Our *ex vivo* microbiome was highly impacted by familial relationship (Table 6, Fig 4). Further, mice differed in the amount of *Clostridium* cultured. Mouse 1 and 2 had higher numbers of *Clostridium* than mouse 3 and 4 (Fig 7F). Not only were these mice siblings but also weaned by different mothers. The effects of weaning are similar to Bian et al. wherein the abundance of an unclassified strain of *Clostridiaceae* was affected by the nursing mother [63]. Our results reiterate the impact of the mother on the microbiome, but also show this dynamic transfers even when explanted from the source.

Essentially, our results indicate that the feces and large distal colon are highly similar; therefore, it is reasonable to consider avoiding mouse sacrifice by culturing feces. Future experiments will need to control pH shifts to avoid media-related population dynamics. Since none of the plates maintained the original baseline pH or even homogenate pH, we assume that additional buffering capacity or equilibrating media and culturing in a CO_2_ incubator may create a more stable explanted microbiome, possibly maintaining diversity more similar to *in vivo* microbiota. Even with the pH swings seen here, we were able to culture bacteria that are difficult to routinely culture. Ultimately, this study found that pH was a stronger influencer of community composition than oxygen. pH has a strong influence on the establishment of the microbes that will populate the explant culture. Future efforts at establishing an *ex vivo* mouse microbiome should include additional measures geared towards stabilizing pH in order to avoid community shifts related to physical changes in growth media.

## Acknowledgements

We would like to thank the staff at Texas A&M Agrilife Research and Extension Center for aiding in the project, the Tarleton State University Office of Student Research and Creative Activities and the College of Science and Technology for funding this research, and the Tarleton State University College of Graduate Studies for the assistantship for Mr. Castaneda.

## References

1. Luke K Ursell, Jessica L Metcalf, Laura Wegener Parfrey and RK. Definig the Human Microbiome. NIH Manuscripts. 2013;70: 1–12. doi:10.1111/j.1753-4887.2012.00493.x.Defining

2. Tourneur E, Chassin C. Neonatal immune adaptation of the gut and its role during infections. Clin Dev Immunol. 2013;2013: 1–17. doi:10.1155/2013/270301

3. Schwabe R and JC. The microbiome and cancer. Nat Rev Cancer. 2013;13: 800–812. doi:10.1038/nrc3610

4. Sommer F, Bäckhed F. The gut microbiota — masters of host development and physiology. Nat Publ Gr. 2013;11. doi:10.1038/nrmicro2974

5. Cho I, Blaser MJ. The human microbiome: At the interface of health and disease. Nat Rev Genet. 2012;13: 260–270. doi:10.1038/nrg3182

6. Li Q, Han Y, Dy ABC, Hagerman RJ. The Gut Microbiota and Autism Spectrum Disorders. Front Cell Neurosci. 2017;11. doi:10.3389/fncel.2017.00120

7. Turnbaugh PJ, Bäckhed F, Fulton L, Gordon JI. Diet-Induced Obesity Is Linked to Marked but Reversible Alterations in the Mouse Distal Gut Microbiome. Cell Host Microbe. 2008;3: 213–223. doi:10.1016/j.chom.2008.02.015

8. Ericsson AC, Franklin CL. Manipulating the gut microbiota: Methods and challenges. ILAR J. 2015;56: 205–217. doi:10.1093/ilar/ilv021

9. Böhm L, Torsin S, Tint SH, Eckstein MT, Ludwig T, Pérez JC. The yeast form of the fungus Candida albicans promotes persistence in the gut of gnotobiotic mice. PLoS Pathog. 2017;13: 1–26. doi:10.1371/journal.ppat.1006699

10. Hart ML, Ericsson AC, Lloyd KCK, Grimsrud KN, Rogala AR, Godfrey VL, et al. Development of outbred CD1 mouse colonies with distinct standardized gut microbiota profiles for use in complex microbiota targeted studies. Sci Rep. 2018;8: 1–11. doi:10.1038/s41598-018-28448-0

11. Roeselers G, Ponomarenko M, Lukovac S, Wortelboer HM. Ex vivo systems to study host–microbiota interactions in the gastrointestinal tract. Best Pract Res Clin Gastroenterol. 2013;27: 101–113. doi:https://doi.org/10.1016/j.bpg.2013.03.018

12. Lagier JC, Khelaifia S, Alou MT, Ndongo S, Dione N, Hugon P, et al. Culture of previously uncultured members of the human gut microbiota by culturomics. Nat Microbiol. 2016;1. doi:10.1038/nmicrobiol.2016.203

13. Browne HP, Forster SC, Anonye BO, Kumar N, Neville BA, Stares MD, et al. Culturing of “unculturable” human microbiota reveals novel taxa and extensive sporulation. Nature. Nature Publishing Group; 2016;533: 543–546. doi:10.1038/nature17645

14. Rodriguez-Palacios A, Aladyshkina N, Ezeji JC, Erkkila HL, Conger M, Ward J, et al. “Cyclical bias” in microbiome research revealed by a portable germ-free housing system using nested isolation. Sci Rep. Springer US; 2018;8: 1–18. doi:10.1038/s41598-018-20742-1

15. Trosvik P, de Muinck EJ. Ecology of bacteria in the human gastrointestinal tract--identification of keystone and foundation taxa. Microbiome. 2015;3: 44. doi:10.1186/s40168-015-0107-4

16. Goodman AL, Kallstrom G, Faith JJ, Reyes A, Moore A, Dantas G, et al. Extensive personal human gut microbiota culture collections characterized and manipulated in gnotobiotic mice. Proc Natl Acad Sci. 2011;108: 6252–6257. doi:10.1073/pnas.1102938108

17. Shokralla S, Spall JL, Gibson JF, Hajibabaei M. Next-generation sequencing technologies for environmental DNA research. Mol Ecol. 2012;21: 1794–1805. doi:10.1111/j.1365-294X.2012.05538.x

18. Zheng L, Kelly CJ, Colgan SP. Physiologic hypoxia and oxygen homeostasis in the healthy intestine. A Review in the Theme: Cellular Responses to Hypoxia. Am J Physiol - Cell Physiol. 2015;309: C350–C360. doi:10.1152/ajpcell.00191.2015

19. Brady JA, Faske JB, Castañeda-Gill JM, King JL, Mitchell FL. High-throughput DNA isolation method for detection of Xylella fastidiosa in plant and insect samples. J Microbiol Methods. Elsevier B.V.; 2011;86: 310–312. doi:10.1016/j.mimet.2011.06.007

20. Herlemann DPR, Labrenz M, Jürgens K, Bertilsson S, Waniek JJ, Andersson AF. Transitions in bacterial communities along the 2000 km salinity gradient of the Baltic Sea. ISME J. 2011;5: 1571–1579. doi:10.1038/ismej.2011.41

21. Klindworth A, Pruesse E, Schweer T, Peplies J, Quast C, Horn M, et al. Evaluation of general 16S ribosomal RNA gene PCR primers for classical and next-generation sequencing-based diversity studies. Nucleic Acids Res. 2013;41: 1–11. doi:10.1093/nar/gks808

22. Illumina. 16S Metagenomic Sequencing Library Preparation. Illumina.com. 2013; 1–28. Available: http://support.illumina.com/content/dam/illumina-support/documents/documentation/chemistry_documentation/16s/16s-metagenomic-library-prep-guide-15044223-b.pdf

23. Caporaso JG, Lauber CL, Walters WA, Berg-Lyons D, Lozupone CA, Turnbaugh PJ, et al. Global patterns of 16S rRNA diversity at a depth of millions of sequences per sample. Proc Natl Acad Sci. 2011;108: 4516–4522. doi:10.1073/pnas.1000080107

24. Edgar RC. Search and clustering orders of magnitude faster than BLAST. Bioinformatics. 2010;26: 2460–2461. doi:10.1093/bioinformatics/btq461

25. DeSantis TZ, Hugenholtz P, Larsen N, Rojas M, Brodie EL, Keller K, et al. Greengenes, a chimera-checked 16S rRNA gene database and workbench compatible with ARB. Appl Environ Microbiol. 2006;72: 5069–5072. doi:10.1128/AEM.03006-05

26. Cole JR, Wang Q, Cardenas E, Fish J, Chai B, Farris RJ, et al. The Ribosomal Database Project: Improved alignments and new tools for rRNA analysis. Nucleic Acids Res. 2009;37: 141–145. doi:10.1093/nar/gkn879

27. Paulson JN, Colin Stine O, Bravo HC, Pop M. Differential abundance analysis for microbial marker-gene surveys. Nat Methods. 2013;10: 1200–1202. doi:10.1038/nmeth.2658

28. Caporaso JG, Kuczynski J, Stombaugh J, Bittinger K, Bushman FD, Costello EK, et al. QIIME allows analysis of high-throughput community sequencing data. Nat Methods. Nature Publishing Group; 2010;7: 335. Available: http://dx.doi.org/10.1038/nmeth.f.303

29. R Core Team. R: A Language and Environment for Statistical Computing [Internet]. Vienna, Austria; 2018. Available: https://www.r-project.org/

30. McMurdie PJ, Holmes S. Phyloseq: An R Package for Reproducible Interactive Analysis and Graphics of Microbiome Census Data. PLoS One. 2013;8. doi:10.1371/journal.pone.0061217

31. Wickham H. ggplot2: Elegant Graphics for Data Analysis [Internet]. Springer-Verlag New York; 2016. Available: http://ggplot2.org

32. Oksanen J, Kindt R, Legendre P, O’Hara B, Simpson GL, Solymos P, et al. vegan: Community Ecology Package [Internet]. 2008. Available: http://cran.r-project.org/,

33. Yap BW, Sim CH. Comparisons of various types of normality tests. J Stat Comput Simul. 2011;81: 2141–2155. doi:10.1080/00949655.2010.520163

34. Dinno A. dunn.test: Dunn’s Test of Multiple Comparisons Using Rank Sums [Internet]. 2017. Available: https://cran.r-project.org/package=dunn.test

35. Clarke KR, Gorley RN. PRIMER v7. 2015;

36. Faust K, Raes J. CoNet app: inference of biological association networks using Cytoscape [version 1; referees : 2 approved with reservations] Referee Status: F1000Research. 2017;1519: 1–14. doi:10.12688/f1000research.9050.1

37. Shannon P, Markiel A, Owen Ozier 2, Baliga NS, Wang JT, Ramage D, et al. Cytoscape: a software environment for integrated models of biomolecular interaction networks. Genome Res. 2003; 2498–2504. doi:10.1101/gr.1239303.metabolite

38. Barberán A, Bates ST, Casamayor EO, Fierer N. Using network analysis to explore co-occurrence patterns in soil microbial communities. ISME J. 2012;6: 343–351. doi:10.1038/ismej.2011.119

39. Faust K, Sathirapongsasuti JF, Izard J, Segata N, Gevers D, Raes J, et al. Microbial co-occurrence relationships in the Human Microbiome. PLoS Comput Biol. 2012;8. doi:10.1371/journal.pcbi.1002606

40. Roggenbuck M, Bærholm Schnell I, Blom N, Bærlum J, Bertelsen MF, Pontén TS, et al. The microbiome of New World vultures. Nat Commun. 2014;5: 1–8. doi:10.1038/ncomms6498

41. Weldon L, Abolins S, Lenzi L, Bourne C, Riley EM, Viney M. The gut microbiota of wild mice. PLoS One. 2015;10: 1–15. doi:10.1371/journal.pone.0134643

42. Kreisinger J, Čížková D, Vohánka J, Piálek J. Gastrointestinal microbiota of wild and inbred individuals of two house mouse subspecies assessed using high-throughput parallel pyrosequencing. Mol Ecol. 2014;23: 5048–5060. doi:10.1111/mec.12909

43. Ormerod KL, Wood DLA, Lachner N, Gellatly SL, Daly JN, Parsons JD, et al. Genomic characterization of the uncultured Bacteroidales family S24-7 inhabiting the guts of homeothermic animals. Microbiome. Microbiome; 2016;4: 1–17. doi:10.1186/s40168-016-0181-2

44. Lagier JC, Edouard S, Pagnier I, Mediannikov O, Drancourt M, Raoult D. Current and past strategies for bacterial culture in clinical microbiology. Clin Microbiol Rev. 2015;28: 208–236. doi:10.1128/CMR.00110-14

45. Khelaifia S, Raoult D, Drancourt M. A Versatile Medium for Cultivating Methanogenic Archaea. PLoS One. 2013;8. doi:10.1371/journal.pone.0061563

46. Meng J, Xu J, Qin D, He Y, Xiao X, Wang F. Genetic and functional properties of uncultivated MCG archaea assessed by metagenome and gene expression analyses. ISME J. Nature Publishing Group; 2014;8: 650–659. doi:10.1038/ismej.2013.174

47. Gorlas A, Robert C, Gimenez G, Drancourt M, Raoult D. Complete genome sequence of Methanomassiliicoccus luminyensis, the largest genome of a human-associated Archaea species. Journal of Bacteriology. 2012. p. 4745. doi:10.1128/JB.00956-12

48. Peterfreund GL, Vandivier LE, Sinha R, Marozsan AJ, Olson WC, Zhu J, et al. Succession in the Gut Microbiome following Antibiotic and Antibody Therapies for Clostridium difficile. PLoS One. 2012;7. doi:10.1371/journal.pone.0046966

49. Zitomersky NL, Atkinson BJ, Franklin SW, Mitchell PD, Snapper SB, Comstock LE, et al. Characterization of Adherent Bacteroidales from Intestinal Biopsies of Children and Young Adults with Inflammatory Bowel Disease. PLoS One. 2013;8. doi:10.1371/journal.pone.0063686

50. Fisher K, Phillips C. The ecology, epidemiology and virulence of Enterococcus. Microbiology. 2009;155: 1749–1757. doi:10.1099/mic.0.026385-0

51. Winstedt L, Frankenberg L, Hederstedt L, Von Wachenfeldt C. Enterococcus faecalis V583 contains a cytochrome bd-type respiratory oxidase. J Bacteriol. 2000;182: 3863–3866. doi:10.1128/JB.182.13.3863-3866.2000

52. Ramsey M, Hartke A, Huycke MM. The Physiology and Metabolism of Enterococci - PubMed - NCBI. Enterococci: From Commensals to Leading Causes of Drug Resistant Infection. 2014. pp. 424–465. Available: http://www.ncbi.nlm.nih.gov/pubmed/24649507%5Cnhttp://www.ncbi.nlm.nih.gov/books/NBK190424/

53. Houghtelling PD. Why is initial bacterial colonization of the intestine important to the infant’s and child’s health? J Pediatr Gastroenterol Nutr. 2016;60: 294–307. doi:10.1097/MPG.0000000000000597.Why

54. Wampach L, Heintz-Buschart A, Hogan A, Muller EEL, Narayanasamy S, Laczny CC, et al. Colonization and succession within the human gut microbiome by archaea, bacteria, and microeukaryotes during the first year of life. Front Microbiol. 2017;8: 1–21. doi:10.3389/fmicb.2017.00738

55. Smith EA, Macfarlane GT. Studies on amine production in the human colon: Enumeration of amine forming bacteria and physiological effects of carbohydrate and pH. Anaerobe. 1996;2: 285–297. doi:10.1006/anae.1996.0037

56. Busse HJ. Polyamines. Methods Microbiol. 2011;38: 239–259. doi:10.1016/B978-0-12-387730-7.00011-5

57. Doré J, Blottière H. The influence of diet on the gut microbiota and its consequences for health. Curr Opin Biotechnol. Elsevier Ltd; 2015;32: 195–199. doi:10.1016/j.copbio.2015.01.002

58. Gänzle MG, Follador R. Metabolism of oligosaccharides and starch in lactobacilli: A review. Frontiers in Microbiology. 2012. pp. 1–15. doi:10.3389/fmicb.2012.00340

59. Ilhan ZE, Marcus AK, Kang D, Rittmann BE. pH-Mediated Microbial and Metabolic. mSphere. 2017;2: 1–12. doi:10.1128/mSphere

60. Raymann K, Moeller AH, Goodman AL, Ochman H. Unexplored Archaeal Diversity in the Great Ape Gut Microbiome. mSphere. 2017;2: e00026–17. doi:10.1128/mSphere.00026-17

61. Ueda K, Tagami Y, Kamihara Y, Shiratori H, Takano H, Beppu T. Isolation of bacteria whose growth is dependent on high levels of CO2and implications of their potential diversity. Appl Environ Microbiol. 2008;74: 4535–4538. doi:10.1128/AEM.00491-08

62. Mueller NT, Bakacs E, Combellick J, Grigoryan Z, Maria G. The infant microbiome development: mom matters. Trends Mol Med. 2015;21: 109–117. doi:10.1016/j.molmed.2014.12.002.The

63. Bian G, Ma S, Zhu Z, Su Y, Zoetendal EG, Mackie R, et al. Age, introduction of solid feed and weaning are more important determinants of gut bacterial succession in piglets than breed and nursing mother as revealed by a reciprocal cross-fostering model. Environ Microbiol. 2016;18: 1566–1577. doi:10.1111/1462-2920.13272

